# The Neural Correlates of Working Memory Training in Typically Developing Children – Working Paper

**DOI:** 10.1101/2021.05.21.445110

**Authors:** J. S. Jones, A-L. R. Adlam, A. Benatayallah, F. Milton

## Abstract

Working memory training improves children’s cognitive performance on untrained tasks; however, little is known about the underlying neural mechanisms. We investigated this in 32 typically developing children aged 10-14 years (19 girls and 13 boys; Devon, UK) using a randomized controlled design and multi-modal MRI. Training improved working memory performance and increased intrinsic functional connectivity between the bilateral intraparietal sulci. Furthermore, improvements in working memory were associated with greater recruitment of the left middle frontal gyrus on a complex span task. The repeated engagement of fronto-parietal regions during training may increase their activity and functional connectivity over time, affording greater working memory performance. We discuss the plausibility of generalizable cognitive benefits from a neurobiological perspective and implications for neurodevelopmental theory. This is not the version of record.

## Introduction

Working memory, the ability to retain and manipulate information over a short period of time (Baddeley & Hitch, 1994), is considered a core executive function (Miyake et al., 2000), is associated with a wide range of cognitive skills (Barrett et al., 2004), and predicts children’s academic outcomes (Alloway & Alloway, 2010). For these reasons, working memory has been a common target for cognitive training interventions, which are widely used by schools and the public. Meta-analyses have repeatedly shown that training improves performance on untrained working memory tasks (Melby-Lervåg et al., 2016; Sala & Gobet, 2017; Schwaighofer et al., 2015; Shipstead et al., 2012; Soveri et al., 2017). It has been suggested that training, through repeated and prolonged practice on working memory tasks, leads to neuroplastic changes that support increased cognitive capacity (Klingberg, 2010). However, this assumption has not been extensively tested and, on the contrary, evidence for broad improvements in other domains associated with working memory capacity, such as reasoning and academic attainment, has been limited in reviews and meta-analyses (Aksayli et al., 2019; Sala & Gobet, 2019, 2020; Simons et al., 2016). Recent evidence has suggested that working memory training may, more selectively, improve typically developing children’s mathematical ability, at least in the short-term (Jones et al., 2020; Judd & Klingberg, 2021). The purpose of this investigation was to examine the structural and functional neural correlates of working memory training in typically developing children using magnetic resonance imaging (MRI).

Despite numerous behavioral studies, reviews, and meta-analyses there is still no consensus regarding the efficacy of working memory training. Understanding whether and how working memory training impacts the brain can provide important insights into which cognitive processes may be improved. For example, the absence of any change in brain structure or function would be inconsistent with the view that training induces neuroplastic changes that lead to generalizable improvements in working memory capacity. Instead, working memory improvements may be explained by the acquisition and refinement of strategies or cognitive routines that are specific to the cognitive processes they were trained on (Gathercole et al., 2019; von Bastian & Oberauer, 2014), such as grouping for serial recall on span tasks (Dunning & Holmes, 2014). However, as these strategies and routines involve different cognitive operations, certain brain regions may be recruited to lesser or greater extents. Indeed, mnemonic strategies have been shown to engage fronto-parietal regions to different extents (Bor & Owen, 2007). Therefore, to ascertain the potential for generalizable cognitive benefits of training, it is critical to examine the effects of brain function in different contexts, such as different tasks or at rest. Similarly, evidence for changes in brain structure would suggest that the cognitive benefits of training have the potential to be long-lasting and generalizable, to the extent that the regions that change support multiple cognitive processes. To further a mechanistic and neurobiological account of how working memory training works we investigate the underlying neurobiological substrates using multiple measures of brain structure and function across task contexts, which may help to elucidate the potential and the limits of working memory training.

Currently, there is a paucity of research investigating the neural mechanisms of working memory training and, to date, only one controlled study has been conducted in typically developing children. This study specifically investigated resting-state functional connectivity in 27 children using magnetoencephalography (MEG; Astle et al., 2015). Cogmed, a commercially available five week training program on 11 span tasks (Klingberg et al., 2005), was found to improve children’s working memory performance and increased functional connectivity between the right fronto-parietal network and left lateral occipital cortex compared to non-adaptive training, which remained at an easy difficulty throughout. In addition, improvements in working memory across both groups correlated with increased connectivity within the dorsal attention network and between the dorsal attention network and left inferior temporal cortex. The authors suggested that the repeated and demanding co-activation of fronto-parietal regions during training may increase functional connectivity between them and afford greater attentional capacity. However, functional connectivity in the dorsal attention network did not significantly differ between the two training groups over time. A trial of 21 children born extremely preterm using fMRI showed some evidence that working memory training increased functional connectivity within the dorsal attention and default mode networks, compared to non-adaptive training (Tseng et al., 2019). However, these results did not survive correction for multiple comparisons and it is unclear whether the neural effects of training are comparable across different child populations as behavioral outcomes vary (Jolles & Crone, 2012; Melby-Lervåg & Hulme, 2013) and are associated with baseline ability (e.g. Judd & Klingberg, 2021; Rennie et al., 2020). Thus, while existing studies highlight a possible role of the dorsal attention network in working memory training, there is currently no statistically reliable evidence that training causally affects children’s functional connectivity within this network.

Beyond possible effects on functional connectivity, working memory training may also affect the recruitment of brain regions when working memory is engaged. However, there are currently no controlled investigations of task-related brain activation in typically developing children and only three studies have been conducted in other child populations, each with methodological limitations. Two pilot studies reported broad changes in fronto-parietal activity when children were engaged in a visual span task: activity increased following Cogmed in adolescents with ADHD (Stevens et al., 2016), whereas it decreased following four hours of training on three span tasks in children born very preterm (Everts et al., 2017). Increased activation following training may suggest greater neuronal recruitment, whereas decreased activation may suggest greater efficiency from a more precise neural response (C. Kelly et al., 2006). However, interpretation of these mixed findings is significantly limited as neither study compared the effects to a control group. The only controlled investigation in children found that Cogmed was not associated with brain activation changes on an n-back task in 18 children born extremely preterm or with extremely low birthweight, compared to non-adaptive training (C. E. Kelly et al., 2020). However, this may be because children in this sample showed no behavioral improvements from training and/or because the task was relatively distal to training (Anderson et al., 2018). Since there is no evidence in typically developing children and the evidence in other child populations has significant shortcomings, it is currently unclear whether working memory training affects how typically developing children’s brains are functionally activated when working memory is engaged.

Meta-analyses of adult working memory training studies have provided more consistent evidence for activation changes in fronto-parietal regions as well as the striatum. A meta-analysis of eight controlled studies reported decreased activity in the right middle frontal gyrus and right inferior parietal lobe, and affected activity in the putamen (Li et al., 2016). Yet this also included studies with brief practice, which are less typical for cognitive training programs and have more often been associated with activation decreases (see Klingberg, 2010). A more recent meta-analysis of 26 studies also reported decreased activation in the right posterior parietal cortex and increased frontal activations (Salmi et al., 2018). Although no direct analysis was performed, activations compared to perceptual-motor training (as a control) and activations associated with longer training periods (>2 weeks) were consistent in the right dorsolateral prefrontal cortex, the right frontal eye field, and the bilateral insula. Thus the longer training periods adopted in commercially available working memory training programs for children, such as Cogmed, may be associated with activation increases, particularly in fronto-parietal regions implicated in working memory.

Finally, it is possible that changes in brain structure may underlie behavioral improvements observed in working memory training. This has yet to be investigated in typically developing children, but one study has examined whether training alters grey matter volume in 48 children born extremely preterm (C. E. Kelly et al., 2020). Children that received Cogmed showed increased volume in the left lateral occipital cortex compared to non-adaptive training, although this did not survive correction for age. The absence of an effect should be treated with caution because there was no behavioral effect of training in this sample, as previously mentioned (Anderson et al., 2018). The findings in adults are similarly equivocal: one study reported broadly increased fronto-parietal volume but only compared to a passive control group (Takeuchi et al., 2013), whereas a study on mental arithmetic training reported reduced fronto-parietal volume compared to an active control group (Takeuchi et al., 2011), and another study reported no significant change compared to an active control group (Metzler-Baddeley et al., 2016). Thus, the possible effects of training on children’s grey matter volume are currently unclear.

In summary, current evidence suggests that working memory training in childhood may be associated with changes in fronto-parietal function. However, controlled investigations of training-related neuroplasticity in typical development are currently limited to one MEG study of functional connectivity (Astle et al., 2015), which is less optimal for network localization relative to fMRI, and where task-related activity and grey matter volume were not investigated. Importantly, cognitive and neural plasticity may be greatest in childhood, and particularly in typically developing children, where larger training effects have been observed (Jolles & Crone, 2012; Melby-Lervåg & Hulme, 2013), possibly when neural systems are less specialized and have the greatest propensity to change (Wass, 2015; Wass et al., 2012). This leaves open questions about how training might affect typically developing children’s brain structure, as measured by grey matter volume, neuronal recruitment when engaged on working memory tasks, and resting-state networks at a higher spatial resolution. Integrating findings across these measures will be critical to developing a mechanistic account of how training works in the brain and, in turn, how it might be expected to support generalizable cognitive benefits.

### Neurodevelopment

Investigating the neural correlates of working memory training has implications beyond training as an intervention and may provide insights about the neurodevelopment of working memory in childhood. Accumulating evidence suggests that specialization of brain function occurs through activity-dependent regional interactions (Johnson, 2011). Mature working memory in adulthood is associated with a bilateral fronto-parietal network including core regions in the dorsolateral prefrontal cortex, middle frontal gyrus, and superior parietal lobe (Rottschy et al., 2012). Brain activity is more distributed in children and reduced in fronto-parietal regions compared to adults (Geier et al., 2009) but as children’s working memory matures, brain activity becomes more localized to core working memory regions (Scherf et al., 2006). Increases in fronto-parietal activity with age are associated with increases in working memory capacity (Klingberg et al., 2002), suggesting that these changes in brain function are important for cognitive development. This specialization is also observed in resting brain function as neighboring regions segregate and functional connectivity within resting-state networks increases with age (Fair et al., 2013; Satterthwaite, Wolf, et al., 2013). These changes in function may be mediated by changes in brain structure; functional specialization in neighboring regions may occur due to a reduction in local grey matter volume through synaptic pruning, whilst integration between remote neural populations may occur through increased myelination in white matter tracts (Johnson, 2011).

While cross-sectional and longitudinal studies have demonstrated general neurodevelopmental trends, which may arise from maturation and/or experience, working memory training provides a relatively unique opportunity to examine the specific effects of cognitive experience in isolation. Indeed, training can be used to experimentally manipulate children’s general working memory performance and to observe causal effects on brain function and structure. Repeatedly activating the fronto-parietal regions that support working memory through training may lead to refinements in this functional network, such as more precise neuronal recruitment or increased coupling within the network, which are also observed with age. It is possible that these experience-dependent effects may mirror those occurring in typical development.

### The Present Study

The first aim of the present study is to investigate how working memory training works by examining a mechanistic, neurobiological account of working memory training in typically developing children using multi-modal MRI. Investigating the effects of training at the level of biological substrates may reveal novel insights into the current lack of consensus on the potential for generalizable cognitive benefits that has arisen from discrepancies in behavioral findings. In turn, this may have wide implications for the current applications of cognitive training in educational and clinical settings. The second aim of the study is to investigate the neurodevelopment of working memory in childhood by isolating the effects of working memory practice from maturation and other experience. To achieve this we will examine whether adaptive Cogmed working memory training in typically developing children is associated with changes in brain activation on a simple and complex span task, resting-state functional connectivity, and grey matter volume, in comparison to non-adaptive control training.

## Method

### Participants

Data are presented for 32 right-handed children (19 girls and 13 boys) that completed the training and final assessments with a mean age of 12 years and 2 months (*SD* = 1.22 years). Left-handed children were not invited to participate to minimize variance at the group-level that may arise from differences in brain lateralization. An additional 18 children did not complete the training and final assessments (see Figure S1 for details of drop outs and missing data). Participants were recruited from Devon, UK, and the majority were White British (95% in similar studies). The final sample included 17 children in the Cogmed adaptive training intervention group and 15 children in the non-adaptive training control group. At baseline, there were no significant differences between the two conditions in IQ, working memory capacity, age, or gender (see Table 1).

**Table 1.** Baseline Characteristics of the Final Sample.

| | Control (n=15) | Cogmed (n=17) | $t(30)$ | $p$ |
| --- | --- | --- | --- | --- |
|  | Mean (SD) | Mean (SD) |  |  |
| **Age (years)** | 11.9 (1.15) | 12.37 (1.27) | 1.1 | 0.281 |
| **IQ** | 116.07 (14.36) | 112 (13.28) | 0.83 | 0.412 |
| **AWMA** | 107 (8.34) | 108.28 (7.43) | 0.46 | 0.649 |

*Note.* Scores on the Dot Matrix and Odd-One-Out indicate proportion of correct responses. Automated Working Memory Assessment (AWMA).

### Design and Procedure

Children and parents/guardians were informed that the aim of the study was to compare two computerized cognitive training programs. Following baseline assessment, 50 children were randomly allocated to receive either adaptive or non-adaptive working memory training in equal numbers (see Figure S1 CONSORT Flow Diagram). Two children were excluded before randomization because they could not tolerate the MRI scanner. Parents/Guardians and children were instructed on how to use their respective training program and they practiced for approximately 10 minutes until they were confident of how they would login and use the program at home. Training support was offered for all participating children by certified Cogmed coaches, who sent weekly emails and discussed any relevant issues. A parent/guardian also agreed to be the child’s training aide, which involved organizing training times, managing rewards, and offering encouragement. Training aides were given guidance on how best to support their child’s training. Children were given instructions for the training tasks and a booklet to timetable their training sessions, record their goals, and acknowledge mutually agreed rewards with their parent/guardian. The booklet also included a training agreement that the child, training aide, and coach were requested to sign. Training was completed for approximately 45 minutes a day, five days a week, for five weeks and all participating children were instructed to complete 20-25 training sessions in accordance with the Cogmed protocol. Children were rewarded with an item of stationery for every five sessions completed and a £20 Amazon voucher on completion of the whole program. At the end of the study, children who were assigned to the control group were offered a free license to use Cogmed in their own time.

### Working Memory Training

Cogmed RoboMemo included a battery of 11 short-term and working memory tasks with visuospatial and verbal stimuli (Klingberg et al., 2005). The tasks required recalling a sequence of spatial locations in order, tracking and recalling a sequence of moving spatial locations or objects, reordering and recalling a sequence of spatial locations, recalling a sequence of digits in reverse order, and recognizing a sequence of letters. Each session lasted approximately 45 minutes with breaks and involved training on eight tasks for approximately 15 trials each. The difficulty of the training tasks adapted on a trial-by-trial basis according to the individual’s performance. The complexity of the stimulus sequence or the number of items to remember would increase following successful responses, whereas they would decrease following incorrect responses. To control for time spent training, the number of trials adapted to an individual’s processing time and current span level, where fewer trials were presented for children training at a higher span and more trials were presented for children training at a lower span. Cogmed included a difficulty level meter, high scores, audio and verbal feedback, and a reward game ‘Robo Racing’, which children could play for a few minutes at the end of a session depending on performance.

Children assigned to non-adaptive working memory training practiced an online verbal updating working memory task (Roberts et al., 2021). This control task was chosen because it shares a number of characteristics with the non-adaptive version of Cogmed, which was discontinued shortly before the commencement of the study. At the beginning of each session, children were first required to memorize a list of seven words. Their memory was then assessed on a cued and free recall test and children could only proceed to the updating training after perfect performance. On each trial of the training task, three words from the original list were presented in three separate boxes for 5s (see Figure 1). Children were then required to update their memory of the words in accordance with two consecutive updating sub-tasks, which were randomly selected without replacement from a choice of three (see Figure 1: d-f). The sub-tasks either required replacement of a target word with one that was one or two words further down the original list (Figure 1: d-e), or replacement of a target word with one from the other boxes (Figure 1: f). In the event that the sub-tasks indicated replacement of a word beyond the end of the list, children were instructed to start again at the beginning of the list. Finally, a word from the original word list was presented in one of the boxes and children were asked if it was the correctly updated word for that box. Children were given 5 seconds to respond ‘Yes’ or ‘No’, before it was marked incorrect. Half of the trials required a ‘Yes’ response and half of the trials required a ‘No’ response. Difficulty was fixed throughout. Children were required to remember three words and had to perform two consecutive updating tasks on each trial.

**Figure 1.**
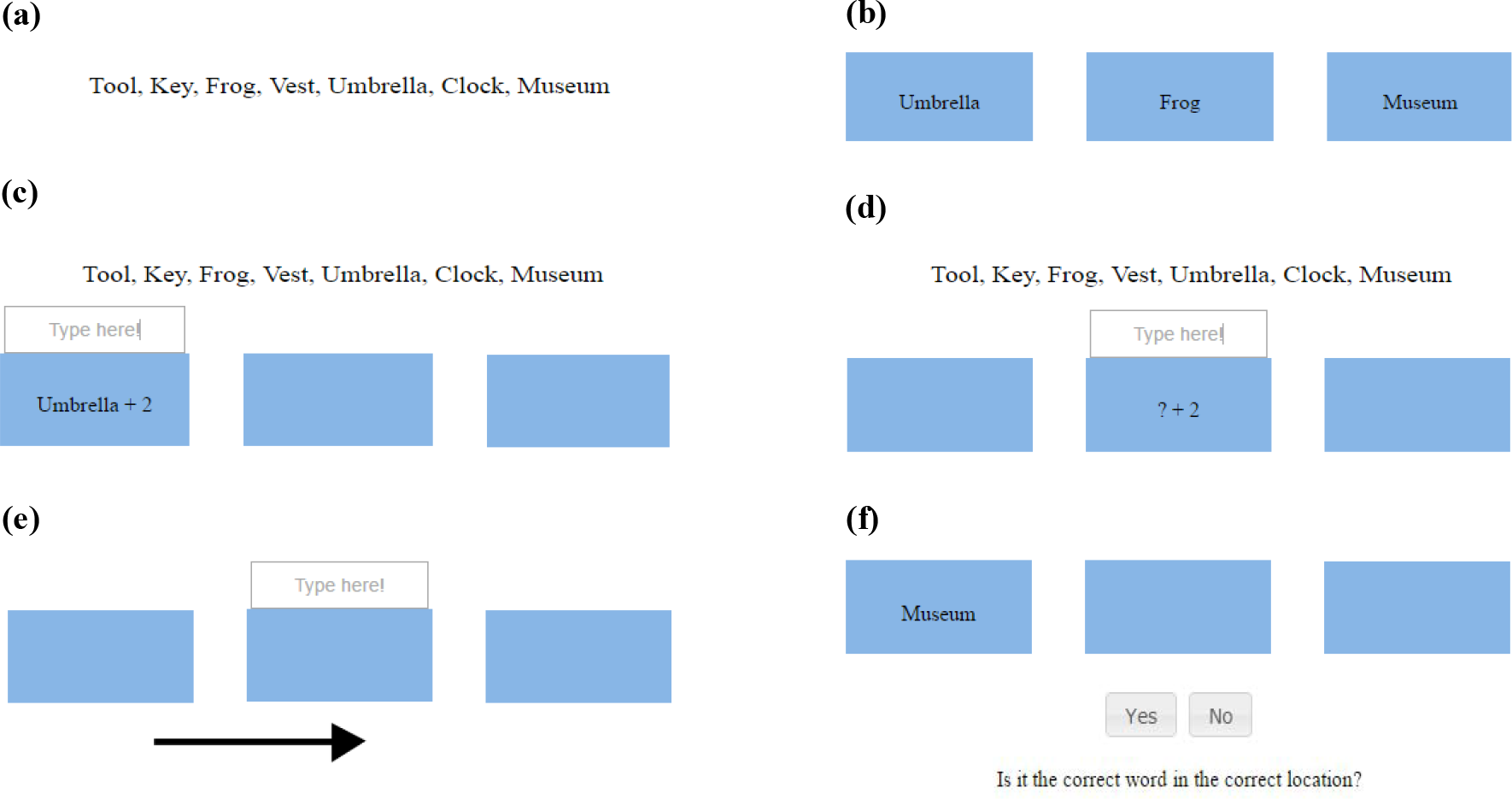
Components of the non-adaptive control updating task. *Note*. (a) a new word list was learnt at the beginning of each training session and was present during the updating sub-tasks; (b) at the start of the trial, three words were presented in Boxes 1-3 (left to right) for 5s; (c) updating sub-task 1 presented ‘+1’ or ‘+2’ next to one of the words, indicating replacement with the word that is one or two forward in the original word list - here the contents of Box 1 (‘Umbrella’) should be replaced with the word that is two forward in the list (‘Museum’); (d) updating sub-task 2 presented a question mark, which masked a word in one of the boxes, and ‘+1’ or ‘+2’, indicating replacement with the word that is one or two forward in the word list - here the contents of Box 2 (‘Frog’) should be replaced with the word that is two forwards (‘Umbrella’); (e) updating sub-task 3 required replacement of the contents of Box 2 (‘Umbrella’) with the contents of Box 1 (‘Museum’), as indicated by the arrow; (f) following two updating sub-tasks, a word was shown in one of the boxes and children decided if it was the correct word.

### Measures

Standardized assessments were administered by trained researchers who were unaware of the child’s group assignment. Working memory was assessed before and after training using eight span tasks from the Automated Working Memory Assessment (AWMA; Alloway, 2007). This included two measures of verbal storage (Digit Recall and Word Recall), two measures of verbal working memory (Backwards Digit Recall and Listening Recall), two measures of visuospatial storage (Mazes Memory and Block Recall), and two measures of visuospatial working memory (Mr. X and Spatial Span). There is good test-retest reliability for these measures ranging from *r* = 0.64-84 (Alloway et al., 2006). There is high agreement between simple and backward/complex span tasks in children (Alloway et al., 2006), suggesting that they measure the same underlying sub-processes (see Unsworth and Engle, 2007). As such, performance across the tasks was averaged for each individual to form an overall composite score of working memory, as in previous studies (e.g., Astle et al., 2015; Jones et al., 2020).

IQ was also assessed to characterize the sample at baseline using the two sub-tests version (FSIQ-2) of the Wechsler Abbreviated Scale of Intelligence-II (WASI-II; Wechsler, 2011). This included a measure of crystallized intelligence (Vocabulary) and a measure of fluid intelligence (Matrix Reasoning). The FSIQ-2 has excellent internal consistency (α = 0.93), test-retest reliability (*r* = 0.87-0.95), and interrater reliability (McCrimmon & Smith, 2013).

### fMRI Tasks

The Dot Matrix was a simple span task (see Figure 2) adapted from the AWMA (Alloway, 2007) and is comparable to tasks used in other neuroimaging investigations of Cogmed in adults (e.g. Brehmer et al., 2011). Four to six red dots were sequentially presented on a 4x4 grid for 900ms each, with no inter-stimulus interval (ISI). The dot locations were pre-randomized, meaning that they were consistent for each participant and each session, and they were never repeated within a trial. After a randomized delay of 1000-3500ms, a probe dot was presented in the grid with a number from one to six within it, indicating the serial order of the probe. Participants were required to indicate if the probe was in the correct location and order as one of the previously presented dots by pressing the left button for ‘yes’ and the right button for ‘no’. The inter-trial interval was randomized between 2000-4000ms. The task consisted of 54 trials across two runs, 18 of each span length (4, 5, and 6). Half of the trials required a correct response and half required an incorrect response, approximately half of the incorrect trials presented lures (n = 13), i.e. probes that were in the same location as a previously presented dot but in a different order.

**Figure 2.**
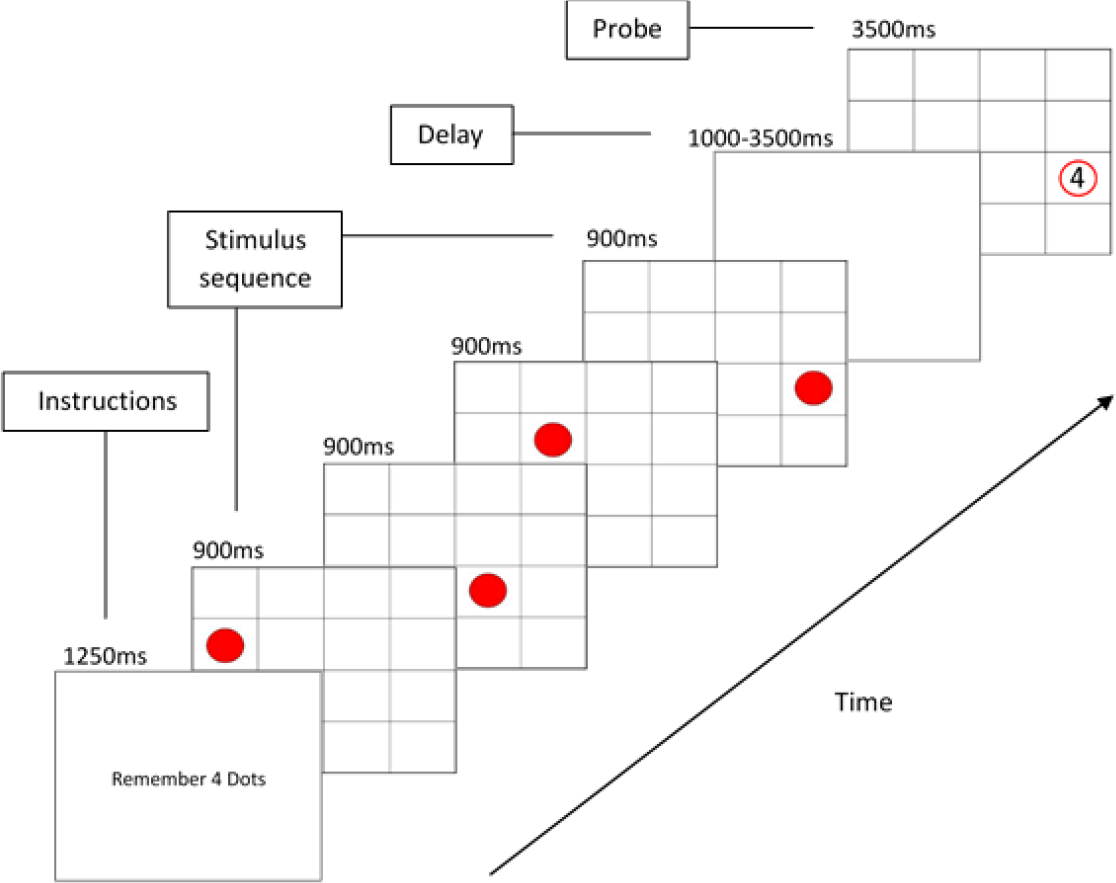
Procedure and timings for the Dot Matrix task. *Note*. An example of a correct probe trial at span four is presented. At the start of each trial, instructions regarding how many stimuli to remember were briefly presented for 1250ms. In the encoding phase, four to six red dots were displayed sequentially on a 4x4 grid for 900ms per stimulus. The stimulus sequence was followed by a randomised delay between 1000 and 3500ms. Finally, a probe was presented for 3500ms and children judged if it was presented in the same location and order as one of the dots from the sequence.

The Odd-One-Out was a complex span task (see Figure 3) also adapted from the AWMA (Alloway, 2007) but has never been used before in published neuroimaging investigations of working memory training. Three to five sets of adjacent shapes were presented sequentially for 2500ms with a 200ms ISI. Each set contained three shapes, two were the same and one was different or ‘odd’. After a 1500ms delay, children were asked to recall the position of one of the odd-one-outs by pressing the appropriate ‘left’, ‘middle’ or ‘right’ button within 4000ms. This was indicated in text by reference to the 1^st^, 2^nd^, 3^rd^, 4^th^, or 5^th^ odd-one-out from the sequence. The inter-trial interval was randomized between 1800-4200ms. The task consisted of 48 trials across three runs, 16 of each span length. Correct responses were equally distributed across the three locations. Both fMRI tasks were programmed in E-Prime 2.0 (Schneider et al., 2002).

**Figure 3.**
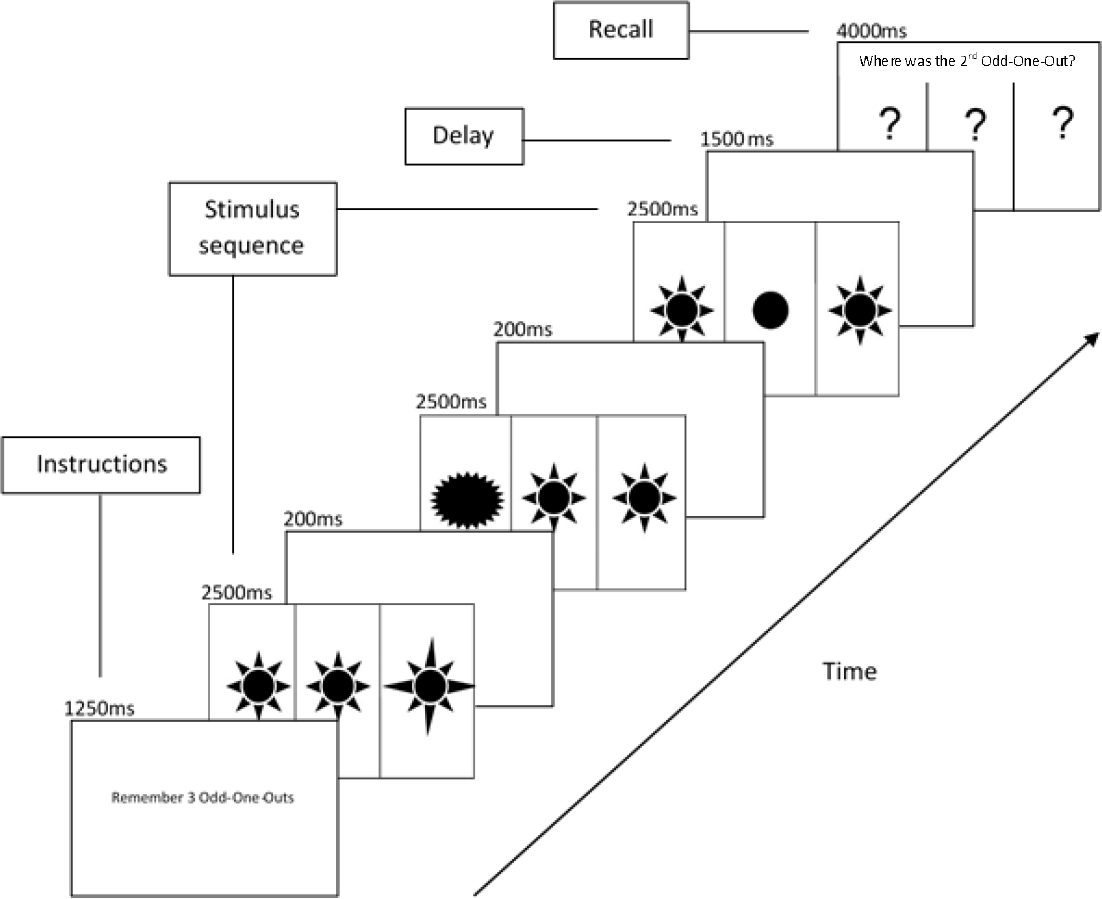
Procedure and timings for the Odd-One-Out task. *Note*. An example of a trial at span three is presented requiring retrieval of the position of the first stimulus. At the start of each trial, instructions regarding how many stimuli to remember were briefly presented for 1250ms. In the encoding phase, three to five stimuli were presented for 2500ms each with a 200ms ISI. Each stimulus consisted of three adjacent shapes; two were identical and one was different, i.e. the odd-one-out. The stimulus sequence was followed by a 1500ms delay. Finally, children had 4000ms to recall the position of one of the odd-one-outs presented in the sequence. The correct response to the example is ‘left’.

### MRI Acquisition

Functional images were acquired using a 1.5T Phillips Gyroscan magnet, equipped with a Sense coil. A T2*-weighted echo planar sequence was used (TR=3000ms, TE= 45ms, flip angle 90°, 35 transverse slices, 2.5 x 2.5 x 3.5mm). Participants completed one scanning session before training and one after training. Each session included two runs of the Dot Matrix task, three runs of the Odd-One-Out task, one run of resting-state, and a structural scan. One hundred and twenty six volumes were collected for each run of the Dot Matrix, 106 volumes were collected for each run of the Odd-One-Out, and 120 volumes were collected for the resting-state in each session. The standard volumetric anatomical MR image was acquired using a 3D T1-weighted pulse sequence (TR = 25ms, TE = 4.2ms, flip angle = 30°, 0.9 x 0.9 x 0.9mm).

### Task fMRI Activation Analysis

The functional images were analyzed using SPM12 (www.fil.ion.ucl.ac.uk/spm). The images were corrected for acquisition order, realigned to the first volume and resliced to correct for motion artefacts. Spatial normalization was performed by coregistering the mean image created from the realigned images to the structural T1 volume. The images were then spatially normalized into the stereotactic space of the Montreal Neurological Institute (MNI). The spatial transformation was applied to the realigned T2* volumes that were spatially smoothed using a Gaussian kernel of 8mm full-width half maximum. Data were high-pass filtered (128s) to account for low frequency drifts. The Blood-Oxygen-Level Dependent (BOLD) response was modeled by a canonical hemodynamic response function (HRF) and the six head movement parameters were included as covariates. Framewise displacement (FD) was calculated to estimate average head movements during each fMRI sequence and compared between the groups overall and over time (Jenkinson et al., 2002). Participants with excessive head movements (mean FD > 0.5mm; Power et al., 2012; Satterthwaite, Elliott, et al., 2013) or substantial artefacts were excluded from each fMRI analysis in a casewise manner. Data acquired for the encoding phase of correct trials were contrasted with the implicit baseline, which is constructed from the BOLD response to phases of the task unrelated to working memory, including the instructions and inter-trial intervals. First-level linear contrasts of parameter estimates for each voxel before and after training were taken to the second-level and a random effects analysis was performed. Activation over time was contrasted between each training condition.

The bilateral middle frontal gyrus (MFG) and superior parietal lobe (SPL) were selected a priori as Regions of Interest (ROI) from the Automated Anatomical Labelling atlas (AAL) (Tzourio-Mazoyer et al., 2002). These regions have previously been associated with activation changes following working memory training in children (Stevens et al., 2016) and adults (Li et al., 2015). Following identical preprocessing of fMRI data detailed above, we analyzed mean activation within each ROI, clusters of activation within each ROI, and clusters of activation in the whole brain as follows. First, group differences in mean activation change within each ROI were analyzed in the Marsbar toolbox (Brett et al., 2002) using a Bonferroni correction for four multiple comparisons (p < 0.0125). Second, clusters of activation change were examined within an explicit mask of the bilateral MFG & SPL using a cluster-forming height threshold of *p* < 0.005 (uncorrected). Finally, whole brain analyses were conducted using a cluster-forming height threshold of *p* < 0.001 (uncorrected). Activation clusters in the ROIs and whole-brain in normalized MNI space were localized to regions of the AAL atlas (Tzourio-Mazoyer et al., 2002). Statistical significance of the ROI and whole-brain clusters was inferred when the cluster extent exceeded that expected by chance (Family-Wise Error [FWE] *p* < 0.05) in Monte-Carlo nonparametric simulations of images with equal smoothness (3dClustSim, AFNI). FWE-corrected critical cluster sizes were 70 for the ROI mask and 111 for the whole-brain.

### Resting-State Functional Connectivity Analysis

Functional connectivity analysis was completed within Conn, version 18a (Whitfield-Gabrieli & Nieto-Castanon, 2012). Pre-processing of the resting-state fMRI was completed using the default Conn pipeline, which implements the same pre-processing steps in SPM12 as outlined for the task-based fMRI above. Identification of global mean intensity and motion outliers was also performed using an automatic artefact detection tool (http://www.nitrc.org/projects/artifact_detect/). Denoising of physiological and motion artefacts included confound regression of six head movement parameters and their first derivatives, motion/global signal outliers, 10 principal component signals estimated from the white matter and cerebrospinal fluid using the aCompCor method (Behzadi et al., 2007), a linear trend, and a band-pass filter 0.008-0.09Hz.

The ROI analyses of the functional connectivity data were conducted using canonically defined resting-state fronto-parietal networks. Two fronto-parietal networks of interest were selected within Conn, which uses data from the Human Connectome Project (Van Essen et al., 2013); the fronto-parietal network comprised of the bilateral lateral prefrontal cortex and posterior parietal cortex, and the dorsal attention network comprised of the bilateral frontal eye fields and intraparietal sulci. Four ROI to ROI analyses were conducted within each network: the left frontal and ipsilateral parietal region, the right frontal and ipsilateral parietal region, the contralateral frontal regions, and the contralateral parietal regions. A Bonferroni correction for multiple comparisons was applied to the analysis of each network (*p* < 0.0125).

### Grey Matter Volume

Voxel-based morphometry (VBM) was used to examine whether Cogmed was associated with changes in regional grey matter volume, compared to the control group. The T1 structural images were analyzed using the DARTEL package (Ashburner, 2007) in SPM12 (www.fil.ion.ucl.ac.uk/spm). The pre- and post-training images were initially co-registered to produce an average image and a divergence image, taking into account the individual difference in time between the scans. The average images were segmented, and the resulting grey and white matter images were spatially aligned using DARTEL. The template and flow fields from DARTEL were used to spatially normalize the divergence images to MNI space, which were then smoothed using a Gaussian kernel of 10mm full-width half maximum.

As the literature is equivocal regarding the effects of working memory training on grey matter volume no regions of interest were selected a priori. Only exploratory whole-brain analyses were conducted at a significance threshold of *p* < 0.001 (uncorrected) and a minimum of 20 contiguous voxels. Clusters in normalized MNI space were localized to regions of the AAL atlas (Tzourio-Mazoyer et al., 2002).

## Results

### Behavioral Outcomes

Related samples t-tests showed that average scores on the eight working memory tasks significantly increased over time for the Cogmed group, Δ +9.06, *t*(16) = 5.83, *p* < 0.001, and for the Control group, Δ +3.84, *t*(14) = 3.13, *p* = 0.007. A mixed ANOVA of the interaction time (pre-training vs post-training) * group (Cogmed vs Control) showed that working memory scores increased significantly more in the Cogmed group compared to the Control group, *F*(1, 30) = 6.7, *p* = 0.015, η_p_^2^ = 0.188 (see Figure 4).

**Figure 4.**
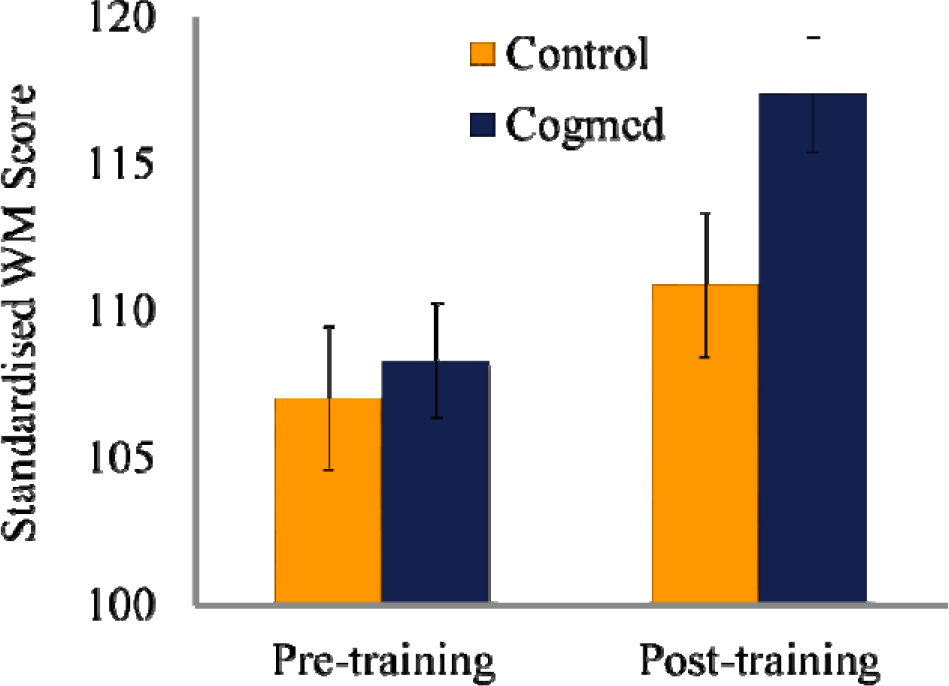
Working Memory Scores Pre- and Post-training for the Two Groups. *Note.* Error bars are 95% confidence intervals with a within-subjects adjustment (Morey, 2008).

### Resting-State Functional Connectivity

Within the dorsal attention network, the Cogmed group showed significantly increased functional connectivity between the left and right intraparietal sulci, relative to the Control group (*p* = 0.005 uncorrected; see Table 2 and Figure 5). Within the lateral fronto-parietal network, the Cogmed group showed no significant changes in functional connectivity relative to the Control group (all *p* > 0.104 uncorrected, see Table S1).

**Figure 5.**
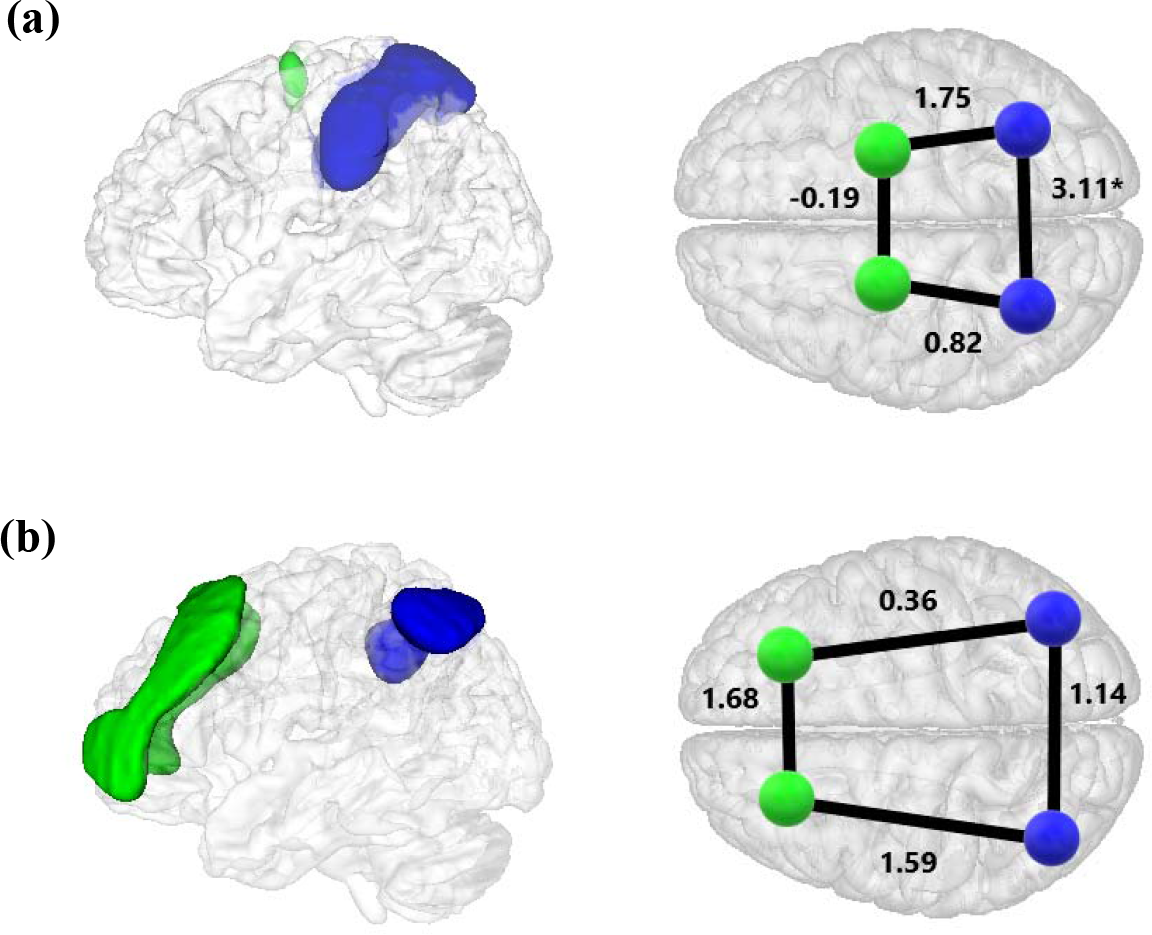
Resting-state network seeds and group comparisons of functional connectivity over time. *Note*. (a) The dorsal attention network consisting of the frontal eye fields and intraparietal sulci and (b) the fronto-parietal network consisting of the lateral prefrontal cortices and inferior parietal lobes. Frontal regions are colored in green and parietal regions in blue. Contrast values of group differences in functional connectivity between the seeds over time are shown on the right where positive values indicate a greater increase in functional connectivity in the Cogmed group. \**p* < 0.0125

**Table 2.**
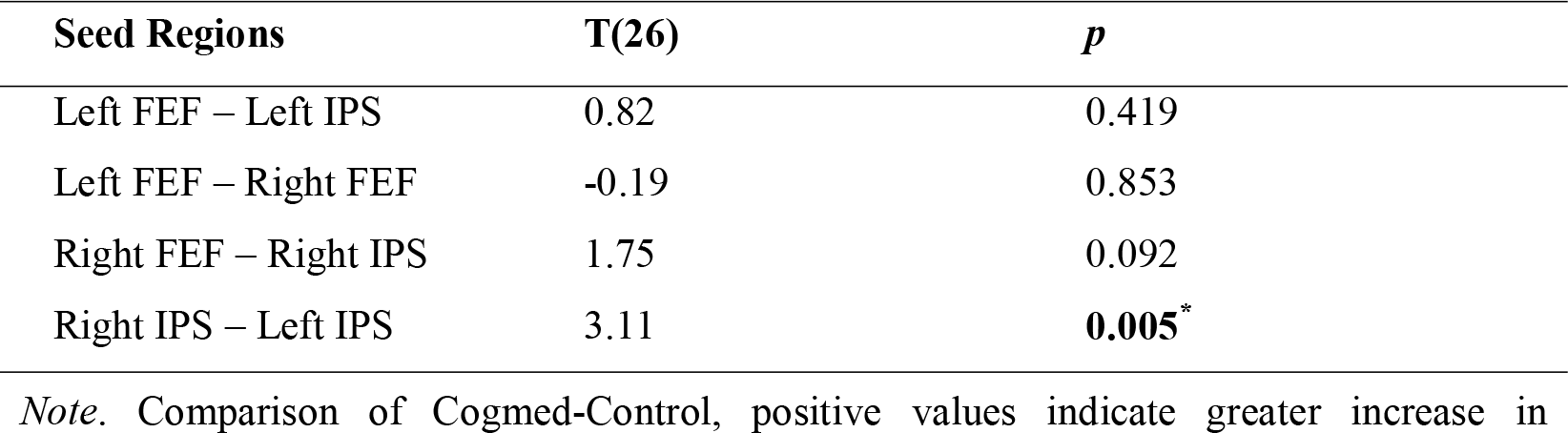
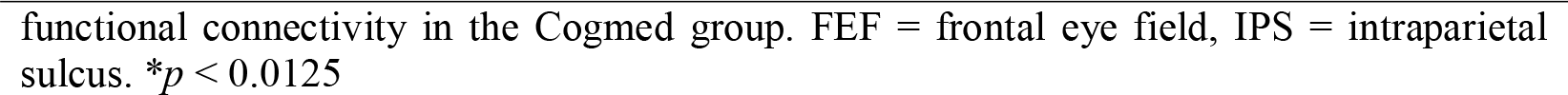
Group comparison of functional connectivity within the dorsal attention network over time.

| Seed Regions | T(26) | *p* |
| --- | --- | --- |
| Left FEF – Left IPS | 0.82 | 0.419 |
| Left FEF – Right FEF | -0.19 | 0.853 |
| Right FEF – Right IPS | 1.75 | 0.092 |
| Right IPS – Left IPS | 3.11 | **0.005\*** |

*Note.* Comparison of Cogmed-Control, positive values indicate greater increase in
functional connectivity in the Cogmed group. FEF = frontal eye field, IPS = intraparietal sulcus. $*p < 0.0125$

### Task-based fMRI

#### Behavior

Behavior on the fMRI tasks is reported for both groups in Table 3. On the Odd-One-Out task, there were no significant group differences in accuracy pre-training, *t*(27) = 1.86, *p* = 0.081, or post-training, *t*(27) = 1.43, *p* = 0.164, there was no main effect of time, *F*(1, 27) = 1.67, *p* = 0.207, and there was no significant group by time interaction, *F*(1, 27) = 0.17, *p* = 0.684. Reaction times did not significantly between the groups pre-training, *t*(27) = 1.4, *p* = 0.173, but the Cogmed group responded significantly faster than the Control group post-training, *t*(27) = 2.73, *p* = 0.011. There was no significant main effect of time, *F*(1, 27) = 1.99, *p* = 0.17, and there was no significant group by time interaction, *F*(1, 27) = 2.18, *p* = 0.151. Thus, while the Cogmed group were faster on the Odd-One-Out post-training, they did not become significantly quicker as a result of training relative to the Control group.

**Table 3.** Behavior on the fMRI tasks during scanning.

|  | Pre-training |  | Post-training |  |
| --- | --- | --- | --- | --- |
|  | Cogmed: M (SD) | Control: M (SD) | Cogmed: M (SD) | Control: M (SD) |
| **Odd-One-Out accuracy** | 0.83 (0.06) | 0.76 (0.14) | 0.85 (0.1) | 0.79 (0.13) |
| **Odd-One-Out reaction time** | 1711.88 (167.6) | 1825.05 (264.89) | 1589.63 (233.44) | 1827.9 (234.19) |
| **Dot Matrix accuracy** | 0.73 (0.09) | 0.71 (0.12) | 0.77 (0.14) | 0.73 (0.15) |
| **Dot Matrix reaction time** | 1559.45 (250.3) | 1612.35 (261.44) | 1440.48 (297.15) | 1519.89 (274.15) |

*Note.* Accuracy on the Dot Matrix and Odd-One-Out indicate proportion of correct responses and reaction times are in milliseconds.

On the Dot Matrix task there were no significant group differences in accuracy pre-training, *t*(27) = 0.5, p = 0.624, or post-training, *t*(27) = 0.79, *p* = 0.434, there was no main effect of time, *F*(1, 27) = 1.66, *p* = 0.209, and there was no significant group by time interaction, *F*(1, 27) = 0.25, *p* = 0.622. There were also no significant group differences in reaction times pre-training, *t*(27) = 0.56, p = 0.582, or post-training, *t*(27) = 0.75, *p* = 0.462, and there was no significant group by time interaction, *F*(1, 27) = 0.15, *p* = 0.699. However, there was a significant main effect of time, *F*(1, 27) = 9.68, *p* = 0.004, indicating that children generally responded faster post-training.

#### Odd-One-Out

Mean activation in the left middle frontal gyrus increased in the Cogmed group relative to the Control group, but this was not significant after correction for multiple comparisons across ROIs (see Table 4).

**Table 4.** Group Comparison of Mean ROI Activation on the Odd-One-Out Task over Time.

| ROI | Contrast | *T*(27) | *p* |
| --- | --- | --- | --- |
| Left Middle Frontal Gyrus | 1.81 | 1.94 | 0.032 |
| Right Middle Frontal Gyrus | 0.46 | 0.47 | 0.321 |
| Left Superior Parietal Lobe | -0.01 | -0.01 | 0.506 |
| Right Superior Parietal Lobe | -0.40 | -0.53 | 0.701 |

*Note.* Positive contrast values indicate a greater increase in mean ROI activation in the Cogmed group compared to the Control group. *p* (uncorrected). \**p* < 0.0125

As both groups performed working memory training, power may have been limited to detect a group difference. Investigating the association between changes in working memory performance and activation change across both groups may be a more sensitive approach (Astle et al., 2015). A regression analysis revealed that improvements in working memory were significantly associated with increased mean activation of the left middle frontal gyrus, *t*(28) = 3.61, *p* < 0.001 (uncorrected, see Table 5 and Figure 6). This was also significant within the Cogmed group, *t*(15) = 2.81, *p* = 0.007 (uncorrected), but not the Control group, *t*(12) = 1.17, *p* = 0.134 (uncorrected; see Tables S5 and S6).

**Figure 6.**
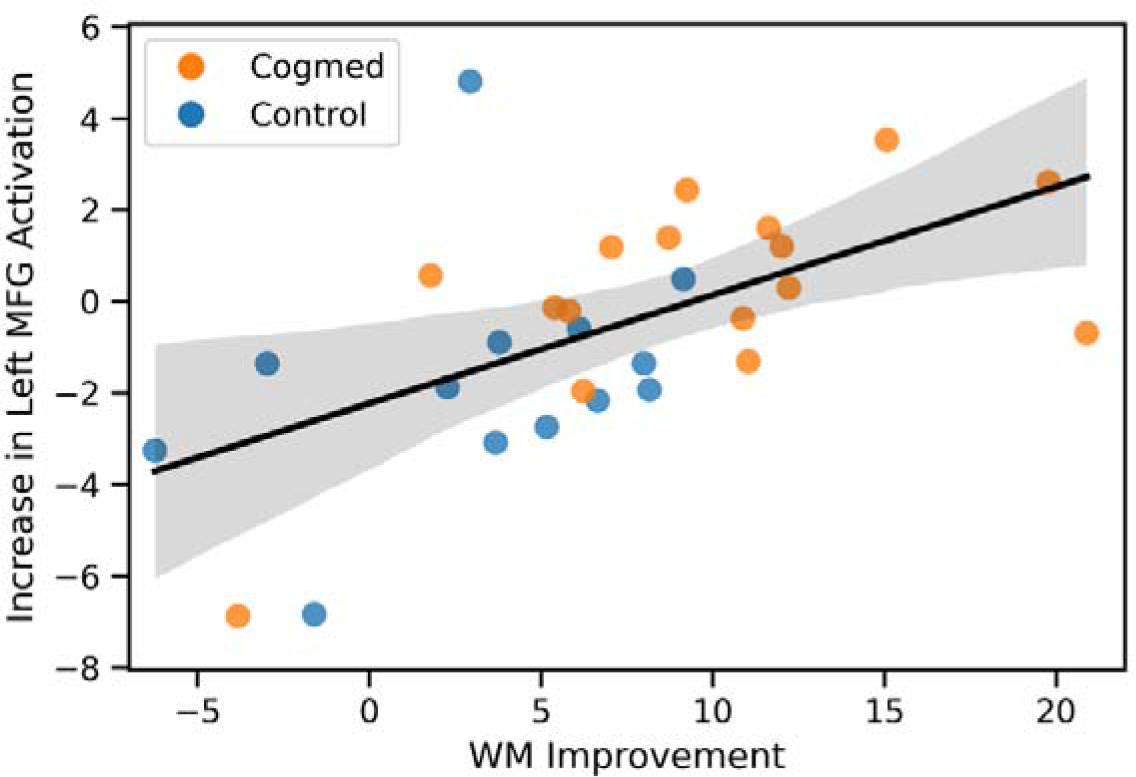
Association between Working Memory Improvements and Left Middle Frontal Gyrus Activation on the Odd-One-Out Task over Time. *Note*. WM = Working Memory, MFG = Middle Frontal Gyrus.

**Table 5.** Association between Working Memory Improvements and Mean ROI Activation on the Odd-One-Out Task over Time.

| ROI | Contrast | *T*(28) | *p* |
| --- | --- | --- | --- |
| Left Middle Frontal Gyrus | 0.24 | 3.61 | < **0.001** * |
| Right Middle Frontal Gyrus | 0.16 | 2.27 | 0.016 |
| Left Superior Parietal Lobe | 0.09 | 1.35 | 0.093 |
| Right Superior Parietal Lobe | 0.07 | 1.27 | 0.107 |

*Note.* Positive contrast values indicate a positive association between mean ROI activation and improvements on the AWMA across both groups. *p* (uncorrected). \**p* < 0.0125

We also examined whether there were significant clusters of voxels that showed a change in activation for the Cogmed group relative to the Control group. Exploratory analyses revealed small clusters of increased activation in the bilateral middle frontal gyri in the Cogmed group, relative to the Control group (see Figure 7 & Table S2). However, no clusters significantly exceeded cluster size expected by chance in simulated images of the ROI mask (k > 70) or whole brain (k > 111; see Table S3 & Figure S2).

**Figure 7.**
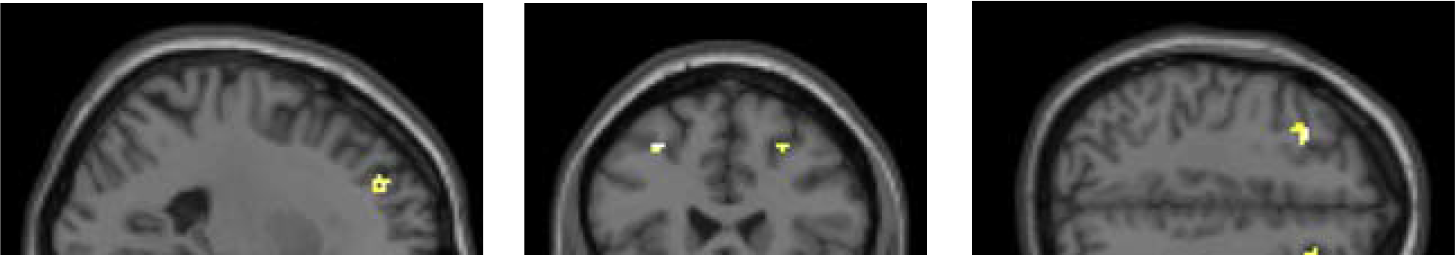
Group by time interaction for the ROI analysis of the Odd-One-Out task.

### Dot Matrix

Cogmed did not increase mean activation in the middle frontal gyri or superior parietal lobes, relative to the Control group (all *p* > 0.419 uncorrected). Furthermore, working memory improvements were not significantly associated with change in mean activation of the ROIs, across both groups (all *p* > 0.093 uncorrected). No significant clusters were detected in the ROI mask (k > 70) or whole brain (k > 111, see Table S4).

### In-Scanner Motion

There were no significant differences in overall head movements between the Cogmed and Control groups (see Table 6). Mixed effects ANOVAs also showed that there were no significant group differences in head movements over time for the resting-state (F(1, 26) = 0.45, p = 0.507), Odd-One-Out (F(1,27) = 2.22, p =0.147), and Dot Matrix (F(1, 27) = 0.24, p = 0.632) tasks.

**Table 6.** Mean framewise displacement in each group.

|  | **Cogmed: M (SD)** | **Control: M (SD)** | ***df*** | ***t*** | ***p*** |
| --- | --- | --- | --- | --- | --- |
| **Resting state** | 0.09 (0.05) | 0.1 (0.05) | 27 | 0.54 | 0.597 |
| **Odd-One-Out** | 0.09 (0.08) | 0.13 (0.09) | 28 | 1.16 | 0.256 |
| **Dot Matrix** | 0.11 (0.07) | 0.16 (0.11) | 28 | 1.68 | 0.104 |

### Grey Matter Volume

There were no significant changes in grey matter volume in the Cogmed group over time, relative to the Control group. Exploratory analyses yielded no clusters greater than 20 contiguous voxels.

## Discussion

We examined the structural and functional neural correlates of working memory training in typically developing children. A significantly larger increase in working memory performance was observed in the adaptive training group (Cogmed), relative to the non-adaptive training group (Control). This was accompanied by changes in brain function but not structure. Specifically, adaptive working memory training significantly increased functional connectivity between the bilateral intraparietal sulci compared to non-adaptive control training. Furthermore, the improvements in working memory after training were significantly associated with greater engagement of the left middle frontal gyrus on a complex span task (the Odd-One-Out).

Consistent with previous evidence, we observed that working memory training was associated with a large increase in typically developing children’s average performance across a range of working memory tasks, relative to control training (Astle et al., 2015; Jones et al., 2020; Sala & Gobet, 2020). The mechanisms of behavioral improvements in working memory as a result of training are still debated. The prolonged mismatch between external demands and current capacity limits during training may induce neuroplastic changes that affords greater capacity (Klingberg, 2010). There is very limited evidence that working memory training broadly improves typically developing children’s cognitive skills associated with working memory that would support this hypothesis (Aksayli et al., 2019; Sala & Gobet, 2019, 2020; Simons et al., 2016); however, recent studies have shown training-related improvements in mathematics (Jones et al., 2020; Judd & Klingberg, 2021), possibly suggesting that far-transfer is more selective. Contemporary evidence in systematic studies of transfer and task similarity increasingly show that transfer is narrow (Gathercole et al., 2019; Rennie et al., 2021), and have suggested that transfer may depend upon the degree to which the training and transfer tasks share the same high-order cognitive routines for effective performance (Gathercole et al., 2019). Neuroimaging can make important contributions to these theories of training-related transfer by identifying neurobiological changes and examining how these contribute to working memory and related cognitive capacities.

Relative to non-adaptive training, we observed that adaptive working memory training significantly increased functional connectivity between the bilateral intra-parietal sulci, which were selected a priori as seed regions of the canonical dorsal attention network. This provides novel evidence that training causally induces functional connectivity changes in regions of this network. Previous studies only showed trend level evidence for a causal effect of training on dorsal attention network connectivity changes relative to a control group, that did not survive correction for multiple comparisons in typically developing children (Astle et al., 2015) or children born extremely preterm (Tseng et al., 2019). This may be because they analyzed average connectivity changes in the dorsal attention network rather than pairwise connectivity changes between core regions of the network, thereby obscuring the specific change in bilateral intraparietal functional connectivity. The dorsal attention network is widely accepted to be involved in the top-down allocation of attention to locations or features (Vossel et al., 2014). Within the network, the intraparietal sulci and frontal eye fields contain retinotopic maps of the contralateral space (Silver & Kastner, 2009), highlighting their significance for visual working memory (Jerde et al., 2012), which was predominantly trained in the Cogmed group. While it is not possible to determine the precise consequences of increased functional connectivity between the bilateral intraparietal sulci on working memory processes in the current study, evidence suggests that it supports manipulation of visuospatial working memory (Bray et al., 2015). Specifically, a previous study showed that maintenance trials preferentially engaged the contralateral intraparietal sulcus and that the ipsilateral intraparietal sulcus was recruited on manipulation trials through interactions with the opposite hemisphere (Bray et al., 2015). It is possible that repeated practice on working memory tasks refined interhemispheric communication between these regions leading to either a more synchronous engagement or a more similar magnitude of response in the contralateral region when manipulating the contents of working memory. Interestingly, bilateral intraparietal functional connectivity has been shown to increase with age and support numerical cognition (Battista et al., 2018; Park et al., 2013), suggesting a plausible neurobiological mechanism for generalizable cognitive benefits of working memory training. Indeed, some of the most promising evidence for far-transfer in typically developing children has been shown in mathematics (Jones et al., 2020; Judd & Klingberg, 2021). Future work will need to determine whether this training-related increase in intraparietal functional connectivity is related to improvements in children’s numerical cognition. Similarly, future work should investigate whether working memory training could ameliorate differences in intraparietal function observed in developmental dyscalculia (Rotzer et al., 2009).

Improvements in working memory after training were also significantly associated with increased activation in the left middle frontal gyrus on a complex span task (the Odd-One-Out). This effect was significant in the adaptive working memory training group but not the non-adaptive control group. To our knowledge, this is the first controlled investigation to demonstrate that improvements in working memory after training are associated with changes in local brain activity in childhood, although we acknowledge that this does not imply causation. Greater increases in left middle frontal activation were observed in the adaptive training group relative to the non-adaptive control group, but this was not significant after a Bonferroni correction for multiple comparisons across the four ROIs. Similarly, the adaptive training group showed greater activation in sub-threshold clusters within the bilateral middle frontal gyri in exploratory analyses. Whilst such a result should obviously be taken with some caution, it does correspond with findings in adults, where a meta-analysis found increased activation of the dorsolateral prefrontal cortex (overlapping with the middle frontal gyrus) in longer interventions (>2 weeks) and compared to a control group (Salmi et al., 2018). Increased middle frontal activation has also been observed in adolescents with and without ADHD after training (Stevens et al., 2016). However, this study reported widespread activation increases and may have been related to test-retest effects because there was no control group. We find that activation increases are much more localized when the active ingredients of working memory training are isolated. A previous controlled study in children born extremely preterm or with extremely low birthweight found no evidence that working memory training affected neuronal recruitment on an n-back task (C. E. Kelly et al., 2020); however, this may be because children showed no cognitive improvements after training (Anderson et al., 2018). Taken together, our findings suggest that improvements in children’s working memory after training are associated with greater engagement of the left middle frontal gyrus, although further work is needed to ascertain whether this effect is causally modulated by the active ingredients of working memory training.

Our findings provide support for functional neuroplasticity after training and when considered with the extant literature show emerging evidence for a neurobiological account of working memory training. The repeated activation of fronto-parietal regions involved in working memory processes during training may increase their activity and connectivity over time thereby affording greater performance on working memory tasks. Indeed, neurofeedback studies have shown that directly manipulating fronto-parietal activity increases functional connectivity within the network and can improve working memory performance (Shen et al., 2015; Zhang et al., 2013, 2016). These training-related changes in brain function over time may be the result of cellular changes in neuronal populations initiated by long-term potentiation (Lynch, 2004). Similar neural mechanisms have been reported for learning spatial routes, whereby hippocampal activity and functional connectivity changed with training (Keller & Just, 2016). This study further demonstrated that training led to short-term changes in hippocampal white matter diffusivity, a measure of structural connectivity, which correlated with behavioral improvements. Although we show no evidence that working memory training alters structure in grey matter volume, largely consistent with other controlled studies in children (C. E. Kelly et al., 2020) and adults (Metzler-Baddeley et al., 2016), there is good evidence that cognitive training alters white matter connectivity from a recent meta-analysis of adult studies (Kristensen et al., 2018). It has been suggested that longer-term changes in myelination through the activity-dependent recruitment of glial cells may support these increases in structural connectivity and could support more stable adaptations to brain function (Kristensen et al., 2018). Similar neuroplastic processes may occur in children’s white matter tracts during working memory training. Increased functional connectivity between the bilateral intraparietal sulci when children were at rest could be the result of more efficient transmission between these regions through increased myelination of the connecting callosal white matter fibers. Future work will need to confirm this hypothesis and the relation between these structural and functional changes over time.

Our findings parallel neurodevelopmental trends in brain function. Functional connectivity typically increases with age between the bilateral intraparietal sulci (Battista et al., 2018; Park et al., 2013) and within resting-state networks, such as the dorsal attention network, more generally (Fair et al., 2013; Satterthwaite, Wolf, et al., 2013). Crucially, while cross-sectional and longitudinal studies have shown the nonspecific effects of age, we demonstrate a causal effect of practice on functional neuroplasticity. Similarly, working memory related activation increases through childhood in core fronto-parietal regions, such as the middle frontal gyrus, and is related to improvements in working memory capacity (Klingberg et al., 2002; Scherf et al., 2006). However, these changes in brain function cannot be attributed to changes in working memory as both correlated with age. We demonstrate that increased recruitment of the middle frontal gyrus is significantly associated with training-related improvements in working memory performance outside the scanner. Importantly, age is unlikely to have substantial effects on brain function or working memory in our study due to the short timescale. This suggests that functional specialization may occur through similar activity-dependent regional interactions (Johnson, 2011) when working memory is engaged either through training or experiences in typical development. For example, schooling places progressive demands on children’s working memory which likely engage fronto-parietal regions and could initiate activity-dependent functional neuroplasticity. Training interventions warrant further investigation as a means to study practice-related neurodevelopment in highly controlled settings.

Concerning strengths and limitations of the current investigation, a major strength of our study was the examination of multiple neural measures. This is particularly informative since there are very few controlled studies of the neural correlates of working memory training in children. In addition, our study included the largest sample (*n* = 28-29) to-date of any neuroimaging investigation of working memory training in typically developing children, to the best of our knowledge. However, it should be noted that this may be considered small in the wider field of developmental psychology. Recruiting larger samples will be an important direction for future work in order to confirm our findings and to examine the neural correlates of far-transfer, as far-transfer effects are typically small (e.g. Judd & Klingberg, 2021). A limitation of the current study is the use of non-adaptive training, which may have been less challenging and/or engaging. In addition, the non-adaptive control task was notably different to the adaptive working memory training, which involved multiple tasks. The task required updating the verbal contents of memory and it lacked some motivational features, such as feedback on performance, high scores, thematic graphics, and audio. Despite these differences, behavioral results were very comparable to studies with matched non-adaptive training (Astle et al., 2015; Dunning et al., 2013) and adaptive training on other tasks (Jones et al., 2020), which may better control for expectancy and motivation effects (Shipstead et al., 2012). A potential limitation of our fMRI design is the use of an implicit baseline rather than a control task. We chose this for practical reasons to reduce scanning time and fatigue effects; however, the resulting activations may be less specific to working memory processes. Finally, we note that the sample had above average IQ and slightly higher than average working memory capacity, which may limit generalizability as baseline ability is known to mediate training effects (e.g. Rennie, Zhang, et al., 2020). Future studies will require larger and more diverse samples to investigate whether neural correlates differ in children with low, average, and high ability.

To conclude, we report novel evidence for functional neuroplasticity in typically developing children following working memory training. Working memory training increased functional connectivity between the bilateral intraparietal sulci of the dorsal attention network and greater working memory improvements after training were associated with increased recruitment of the left middle frontal gyrus on a complex span task. Repeated activation of fronto-parietal regions involved in attentional and executive control during training may increase their activity and connectivity over time enabling greater performance on working memory tasks. Training-related changes in brain function paralleled typical neurodevelopmental trends, suggesting activity-dependent specialization through practice as a potentially common mechanism, and warranting the use of training interventions as highly controlled investigations of neurodevelopmental mechanisms. Finally, we provided a plausible neurobiological account of how training-related changes in bilateral intraparietal functional connectivity could support improvements in numerical cognition. Future work is required to confirm whether training-related changes in brain function can improve numerical cognition, and whether they are associated with ecological outcomes (e.g. performance at school), alterations in structural connectivity, and individual differences.

## Acknowledgements

We would like to thank the ESRC for funding this PhD studentship (Ref: 1490438) in which the work was completed. We would also like to thank all of the children and families that participated in this research. Finally, a massive thank you to the undergraduate students that helped with data collection and supporting families through their training: Zoe Dixon, Bronte Graham, Zoe Moody, and Olivia Muir. Behavioral data is available at https://osf.io/qyjgs/ (DOI: 10.17605/OSF.IO/QYJGS) and neuroimaging data is available on request.

