## Supplementary materials for "The Neural Correlates of Working Memory Training in Typically Developing Children – Working Paper"

#### Child Development

**Figure S1.** *CONSORT Flow Diagram*

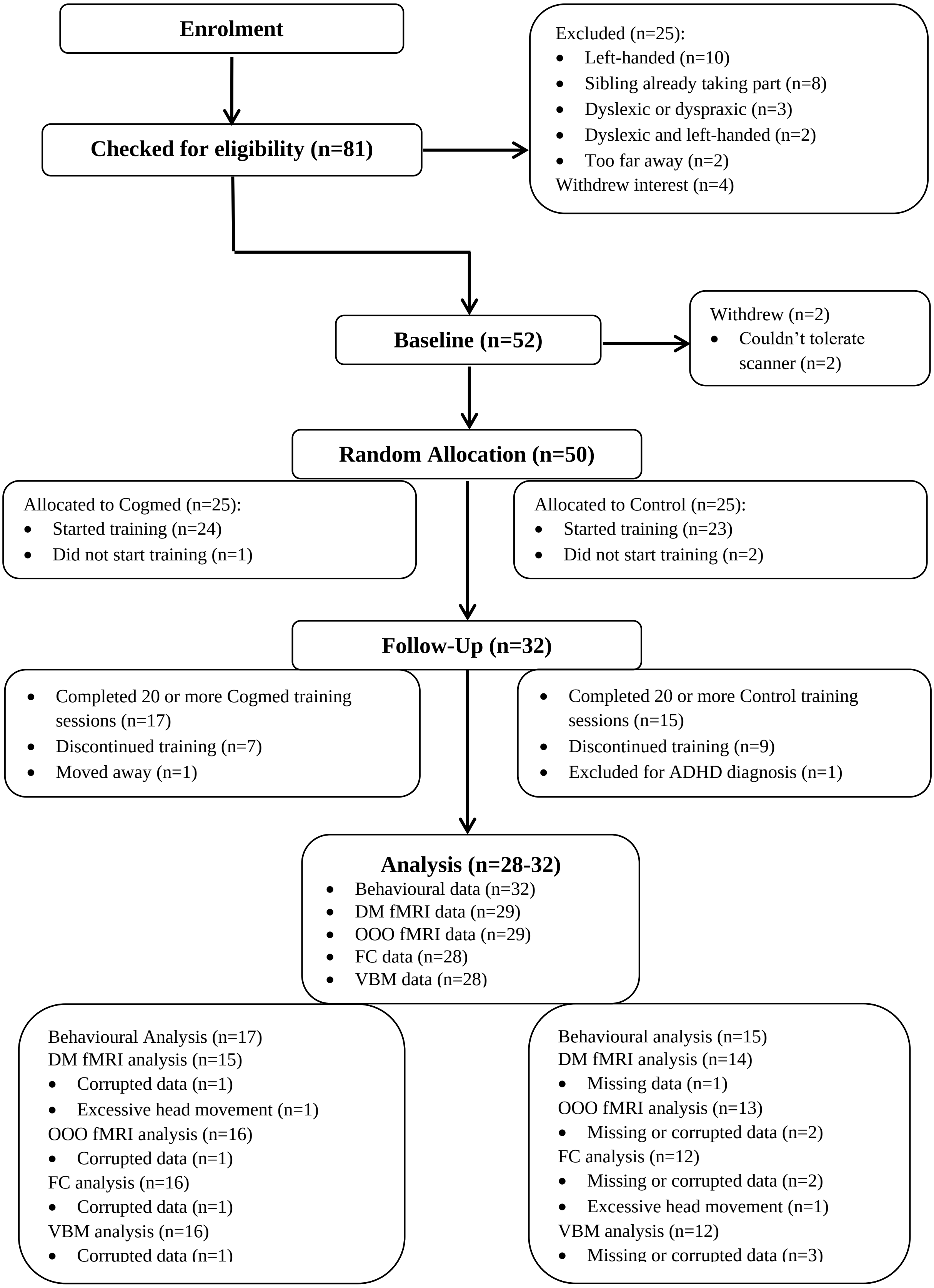


*Note*. DM = Dot Matrix, OOO = Odd-One-Out, FC = Functional Connectivity, VBM = Voxel-Based Morphometry.

### Functional Connectivity in the Lateral Fronto-Parietal Network

Table S1. Group Comparison of Functional Connectivity within the Fronto-Parietal Network over Time.

| **Seed Regions** | ***T*(26)** | ***p*** |
|---|---|---|
| Left LPFC – Left PPC | 1.59 | 0.125 |
| Left LPFC – Right LPFC | 1.68 | 0.105 |
| Right LPFC – Right PPC | 0.36 | 0.721 |
| Right PPC – Left PPC | 1.14 | 0.264 |

*Note.* Comparison of Cogmed-Control, positive values indicate greater increase in functional connectivity in the Cogmed group. LPFC = lateral prefrontal cortex, PPC = posterior parietal cortex. $p$ (uncorrected). $*p < 0.0125$

#### Exploratory Cluster Analysis

Exploratory ROI analysis was conducted within a mask of the bilateral MFG & SPL at a cluster-forming height threshold of $p < 0.005$ and extent of 10 contiguous voxels, as in previous studies (Milton et al., 2012). Exploratory whole-brain analysis was conducted at a cluster-forming height threshold of $p < 0.001$ and extent of 20 contiguous voxels, as in previous studies (e.g. Milton et al., 2012; Milton & Pothos, 2011).

##### Odd-One-Out

The ROI analysis is displayed in Table S1 and Figure S1. Several clusters in the middle frontal gyrus showed greater activation over time in the Cogmed group compared to the Control group, which were localised to BAs 6, 8, and 9. The results of the whole-brain analysis are shown in Table S2 and Figure S2.

Table S2. ROI Group Comparison of Odd-One-Out Task Activation over Time.

| **Brain Region** | **Cluster size** | **Peak *Z*** | **MNI Coordinates** | | |
|---|---|---|---|---|---|
| | | | **x** | **y** | **z** |
| ***Cogmed > Control*** | | | | | |
| Left middle frontal gyrus (BA6/8) | 27 | 3.47 | -32 | 20 | 42 |
| Right middle frontal gyrus (BA8) | 21 | 3.35 | 24 | 28 | 46 |
| Left middle frontal gyrus (BA9) | 23 | 3.18 | -24 | 46 | 26 |
| ***Control > Cogmed*** | | | | | |
| No significant clusters | | | | | |

Table S3. Whole-Brain Group Comparison of Odd-One-Out Task Activation over Time.

| **Brain Region** | **Cluster size** | **Peak Z** | **MNI Coordinates** | | |
|---|---|---|---|---|---|
| | | | **x** | **y** | **z** |
| ***Cogmed > Control*** | | | | | |
| Right posterior cingulate (BA23) | 68 | 4.33 | 16 | -34 | 24 |
| Right parahippocampal gyrus (BA34) | 46 | 3.92 | 26 | 4 | -26 |
| Left superior/medial frontal gyrus (BA6/8) | 69 | 3.87 | -12 | 44 | 40 |
| | | 3.51 | -10 | 30 | 48 |
| | | 3.49 | -6 | 40 | 46 |
| Left anterior cingulate (BA24) | 29 | 3.75 | -18 | -8 | 44 |
| Left amygdala | 23 | 3.48 | -34 | -6 | -22 |
| ***Control > Cogmed*** | | | | | |
| No significant clusters | | | | | |

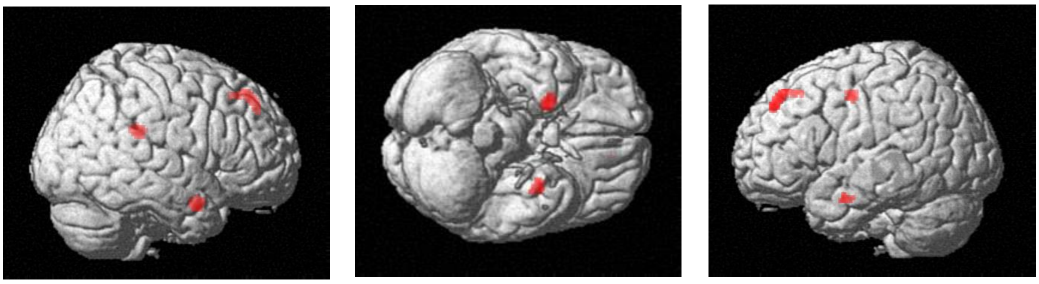

*Figure S3*. Whole brain group comparison of Odd-One-Out activation over time. Regions of increased activation in the Cogmed group relative to control.

#### Dot Matrix

There were no clusters (k >10) in the ROI mask that showed greater activation in the Cogmed group over time compared to the Control group. However, there were clusters in the whole-brain analysis that showed greater and less activation in the Cogmed group over time (see Table S3).

Table S4. Whole Brain Group Comparison of Dot Matrix Task Activation over Time.

| Brain Region | Cluster size | Peak Z | MNI Coordinates | | |
|---|---|---|---|---|---|
| | | | x | y | z |
| ***Cogmed > Control*** | | | | | |
| Left putamen | 22 | 4.00 | -30 | -26 | 0 |
| Left posterior cingulate (BA31) | 44 | 3.94 | -12 | -50 | 28 |
| ***Control > Cogmed*** | | | | | |
| Right parahippocampal gyrus (BA34) | 26 | 4.01 | 14 | -6 | -22 |

#### Working Memory Improvements & ROI Activation on the Odd-One-Out within Groups

Regression analyses examined the association between working memory improvements and changes in ROI activation on the Odd-One-Out task within each group over time. There was a significant association between working memory improvements and left middle frontal activation over time in the Cogmed group (see Table S4), but not for the Control group (see Table S5).

Table S5. Association between WM gain and Mean ROI Activation on the Odd-One-Out Task over Time in the Cogmed group.

| **ROI** | **Contrast** | ***T*(15)** | ***p*** |
|---|---|---|---|
| Left Middle Frontal Gyrus | 0.25 | 2.81 | **0.007*** |
| Right Middle Frontal Gyrus | 0.13 | 1.24 | 0.118 |
| Left Superior Parietal Lobe | 0.17 | 1.54 | 0.073 |
| Right Superior Parietal Lobe | 0.14 | 1.36 | 0.098 |

*Note.* Positive contrast values indicate increased activity over time. $p$ (uncorrected). *$p < 0.0125$

Table S6. Association between WM gain and Mean ROI Activation on the Odd-One-Out Task over Time in the Control group.

| **ROI** | **Contrast** | ***T*(12)** | ***p*** |
|---|---|---|---|
| Left Middle Frontal Gyrus | 0.19 | 1.17 | 0.134 |
| Right Middle Frontal Gyrus | 0.11 | 0.80 | 0.220 |
| Left Superior Parietal Lobe | 0.23 | 1.70 | 0.059 |
| Right Superior Parietal Lobe | 0.08 | 1.25 | 0.118 |

*Note.* Positive contrast values indicate increased activity over time. $p$ (uncorrected). *$p < 0.0125$
